# Design and fabrication of recombinant reflectin-based Bragg reflectors: bio-design engineering and photoisomerism induced wavelength modulation

**DOI:** 10.1101/2020.02.11.942110

**Authors:** Emmanuel Wolde-Michael, Aled D Roberts, Derren J Heyes, Ahu G Dumanli, Jonny J Blaker, Eriko Takano, Nigel S Scrutton

**Affiliations:** EPSRC/BBSRC Future Biomanufacturing Research Hub, Manchester Institute of Biotechnology, Department of Chemistry, The University of Manchester, Manchester, M1 7DN; Department of Materials, The University of Manchester, Manchester, M13 9PL

**Keywords:** reflectins, optically-active thin-films, photoisomerism, Bragg reflectors, angle-dependence

## Abstract

The remarkable camouflage capabilities of cephalopods have inspired many to develop dynamic optical materials which exploit certain design principles and/or material properties from cephalopod dermal cells. Here, the angle-dependent optical properties of various single-layer reflectin thin-films are characterized within the UV-Vis-NIR regions. Following this, the design and fabrication of the first bio-inspired reflectin-based Bragg reflector is described, which was found to conserve the optical properties of single layer films but exhibit a unique characteristic; reduced angle-dependent reflectivity. Finally, a novel method of controlling reflectin thin-film optical properties is introduced; visible light-induced photoisomerism, representing a new class of reflectin-based optical materials.

## INTRODUCTION

Reflectins are a unique family of high-refractive index proteins native to cephalopods (squid, octopus, and cuttlefish).^[1]^ Consisting of repeating motifs (rich in aromatic and sulfur-containing amino acids) separated by positively charged linkers, these proteins are found within cellular Bragg reflectors known as iridophores, where intracellular lamellae containing reflectins are spatially separated by the extracellular matrix (**Figure 1A**).^[2]^ Iridescence arises from coherent Bragg reflection from successive layers, defined by interference theory.^[3]^ Some cephalopods, such as squid in the *Loligidae* family, are able to tightly control the optical properties of these iridophores using a neurotransmitter, acetylcholine (ACh).^[4]^ ACh triggers a signal-transduction cascade which leads to the phosphorylation of reflectins, reducing net charge and triggering reversible hierarchical assembly into a more condensed structure.^[5,6]^ Reduced ion exposure then leads to lamellae dehydration, further increasing the refractive index contrast.^[7]^ Collectively, along with chromatophores (pigment-filled cells which act as spectral filters under neuromuscular control), and leucophores (highly reflective broadband light scatterers), iridophores endow cephalopods with a high level of control over their body coloration and patterning, which they use for both camouflage and signaling.^[1]^ Since their discovery, researchers have been working towards the development of dynamic, optically active biomimetic camouflage technologies which exploit the unique properties of reflectins.^[8–12]^ Facilitating this, recent efforts have been directed toward the characterization of reflectins in vitro, revealing properties such as pH-dependent particle-size formation,^[13]^ the differential roles of motifs and linker regions,^[6]^ the role of key amino-acid residues,^[14]^ reflectin conductive properties,^[15,16]^ and the effect of small molecules on higher-order assembly.^[17]^Advances in the design, fabrication, and characterization of reflectin-based materials have revealed properties such as thickness-dependent coloration,^[9–11,18,19]^ broad near-infra-red (NIR) reflectance,^[11]^ and induced light scattering,^[20,21]^ It has also been demonstrated that the thickness (and therefore optical properties) of these materials can be controlled both pre-fabrication, by varying parameters such as flow-coating angle and sample concentration,^[18,19]^ and post-fabrication, via vapor-induced swelling,^[18,19]^ applying uniaxial strain,^[10]^ or proton conduction.^[9]^ Despite these advancements, the promise of the next-generation of dynamic, tightly-controlled, reflectin-based camouflage technologies is currently stifled by angular-dependent reflectivity, which is a significant yet generally overlooked issue. As most reflectin-based materials are fabricated in the form of thin-films/coatings, their optical reflectivity (*λ*) depends on both the layer thickness (d) and the angle of incidence (viewing angle, *θ*), as shown in equation (1), where n is average refractive index and m is order of reflection. However, while significant efforts have been directed towards developing novel methods of controlling reflectin layer thickness to modulate the thickness-dependent optical properties, little has been done to characterize and/or modify angular-dependence.

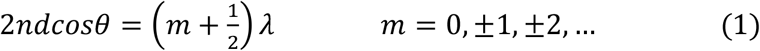

**Figure 1.**
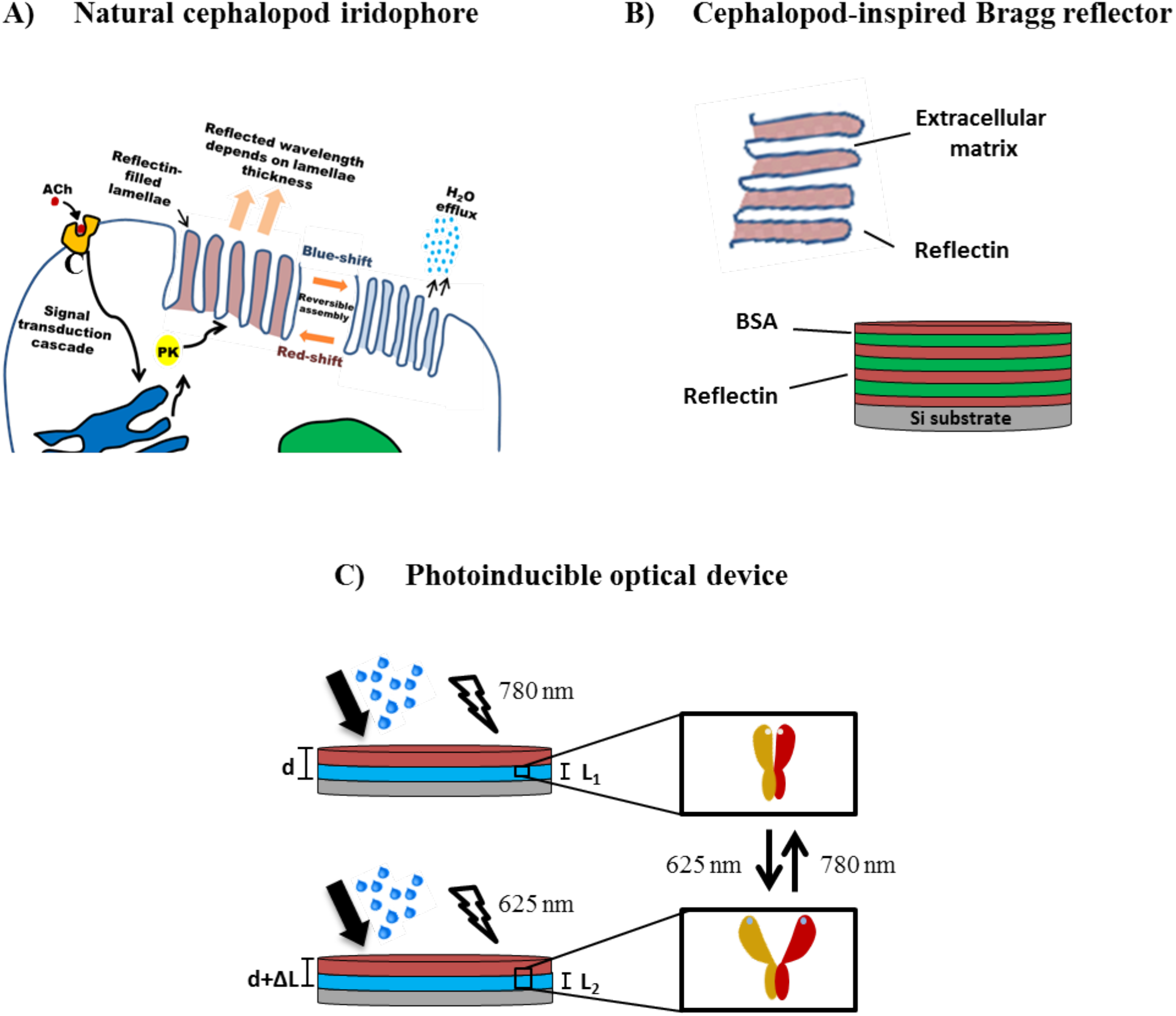
A) Schematic figure showing cephalopod iridophore neurochemical activation. B) Schematic figure of cephalopod-inspired reflectin/BSA Bragg reflector alongside the iridophore Bragg reflector. C) Schematic figure of reflectin-phytochrome device switching between two states. Upon illumination with red/far-red light in the presence of water vapor the phytochrome can convert between an ‘open’ and ‘closed’ state.

## RESULTS

Here, the optical properties of a range of reflectin single-layer thin-films have been characterized over a wide wavelength range (185-3300 nm), revealing previously unreported optical properties. The angular reflectance of both single and multilayered systems was subsequently characterized by varying the angle of incidence between 20 and 70 degrees, revealing significant spectral changes. Following this, the design and fabrication of a cephalopod-inspired reflectin-based Bragg reflector is then outlined (Figure 1B), exhibiting modulated reflectance over a relatively large angle range. Finally, we further introduce a novel method of controlling reflectin thin-film thickness (and therefore optical properties) via visible light-induced isomerism (Figure 1C).

All reflectin isoforms (reflectins are referred to as XXRefYY, where XX is the origin, and YY is the reflectin isoform, Figure S1) were expressed in *Escherichia coli* from recombinant plasmids and were found to be sequestered in inclusion bodies as previously reported.^[13,19]^ Inclusion bodies were isolated using standard inclusion body preparations,^[22]^ solubilized under strongly denaturing conditions, and purified using high pressure liquid chromatography (HPLC) before being lyophilized and stored at 4 °C.^[23]^ Upon spin-coating onto clean Si wafers under the same conditions (Figure 2A), all single-layer films appeared light blue under ambient light upon drying, suggesting comparable thicknesses were achieved (Figure S2).^[3]^ Using cross-sectional scanning electron microscopy (SEM), the thickness of one sample (DORefA2) was determined to be ∼240 nm (Figure 2B, top), in close agreement with atomic force microscopy assessment (AFM, Figure S3). Using equation (1), assuming a refractive index of around ∼1.56,^[24]^ the peak reflectance of a 200 nm single-layer film at normal incidence is around 375 nm, in accordance with the observed colour of our fabricated films. The UV-VIS-NIR reflectance spectra reveal three orders of reflectivity conserved across all isoforms; UV reflectivity at ∼200 nm (2^nd^ order), visible reflectivity at 350-450 nm (1^st^ order), and broad IR reflectivity spanning the near- and short-wave IR region up to 3300 nm (0^th^ order) (Figure S4). Controlling the optical properties in the near- and short-wave IR regions is of particular interest to those in the defense industry as these include commonly surveilled electromagnetic (EM) regions which are often targets for EM signature reduction technologies.^[25]^ By varying the angle of incidence, the angular-dependent reflectivity of these single-layer films was characterized. As the angle of incidence increased from 20 to 70 degrees, the normalized reflectance of all samples at 350-450 nm gradually reduced from ∼1 to ∼0.25 (Figure 2C, top right, Figure S5). Concomitantly, normalized reflectance between 640-750 nm increased from ∼0.25 at 20 degrees to ∼1.5 at 70 degrees. Using equation (1), we can attribute the emergence of this reflectivity at 70 degrees to red-shifting of the 0^th^ order of reflection. Thus, the sustained presence of broad IR reflectivity at these angles suggests that reflectivity in these regions may result from the bulk optical properties of reflectins themselves, rather than thin-film interference. Notably, a peak at ∼2370 nm also emerged at 60/70 degrees, although the significance of this is currently unknown. This angular dependent reflectivity can have negative implications, especially for emerging sensing/anti-camouflage technologies.^[26]^ This therefore represents a major hindrance to the integration of reflectin single-layer films into modern bio-based camouflage technologies. To overcome this seemingly fundamental issue, we began to take more direct inspiration from cephalopods, whose body coloration appears to be angle-independent. With this in mind, more biomimetic configurations of reflectin thin-films were explored to more accurately mimic the structure of iridophores. This led us to design a reflectin-based Bragg reflector consisting of alternating layers of reflectin and bovine serum albumin (BSA), a stable and readily available protein. BSA in this case mimics the presence of extracellular space which separates the multiple reflectin-filled lamellae in iridophores (Figure 1B). Using spin coating, a seven-layered film was fabricated by coating reflectin and BSA sequentially, allowing the film to dry for a few minutes prior to the application of the next layer (Figure 2A). The presence of discrete layers was confirmed using cross-sectional SEM (Figure 2B, bottom), which determined the thickness of a tri-layered reflectin-BSA film to be ∼990 nm. Markedly, while all of the spectral features of our single-layer films were maintained (also confirming the presence of discrete layers of a thickness comparable to that of our single-layer films), angle-dependence in the visible and NIR regions (below 1500 nm) were almost completely reduced (Figure 2C, bottom middle). As the angle of incidence increased from 20 to 70 degrees, normalized reflectance at ∼350-450 nm was maintained around 1, and was retained as the peak reflectance in the visible region. This is the first example of a reflectin-based film whose specular reflectance has been shown to be modulated with respect to viewing angle, and while the presence of a broad reflectance peak may ensure that the impact of angle-dependent reflectivity is minimized,^[10]^ our design does so without compromising spectral coherence. Thus, combined with the dynamic capabilities of the current generation of reflectin-based materials, such as reconfigurable IR reflectivity, ^[10,11]^ moving towards more biomimetic configurations will enable the design of next-generation dynamic, optically active, angle-independent materials. This may represent an important milestone in the reflectin story, enabling a move from developing reflectin-based materials with interesting optical properties to developing reflectin-based camouflage technologies. At wavelengths above 1500 nm, specular reflectance is amplified as angle of incidence increases, with the same peak at around ∼2370 nm emerging at 60/70 degrees, but to an even greater extent. Based on theoretical considerations of multilayer interference,^[3]^ the reduced angular-dependence exhibited by our Bragg reflectors may be considered counterintuitive; however could be explained by the morphology of the coated reflectin, with SEM micrographs revealing the formation of irregular, wrinkled microstructures upon coating (Figure S6). These irregular configurations are thought to arise due to stresses applied during the spin coating process and have been shown to impart unique optical properties.^[27]^

**Figure 2.**
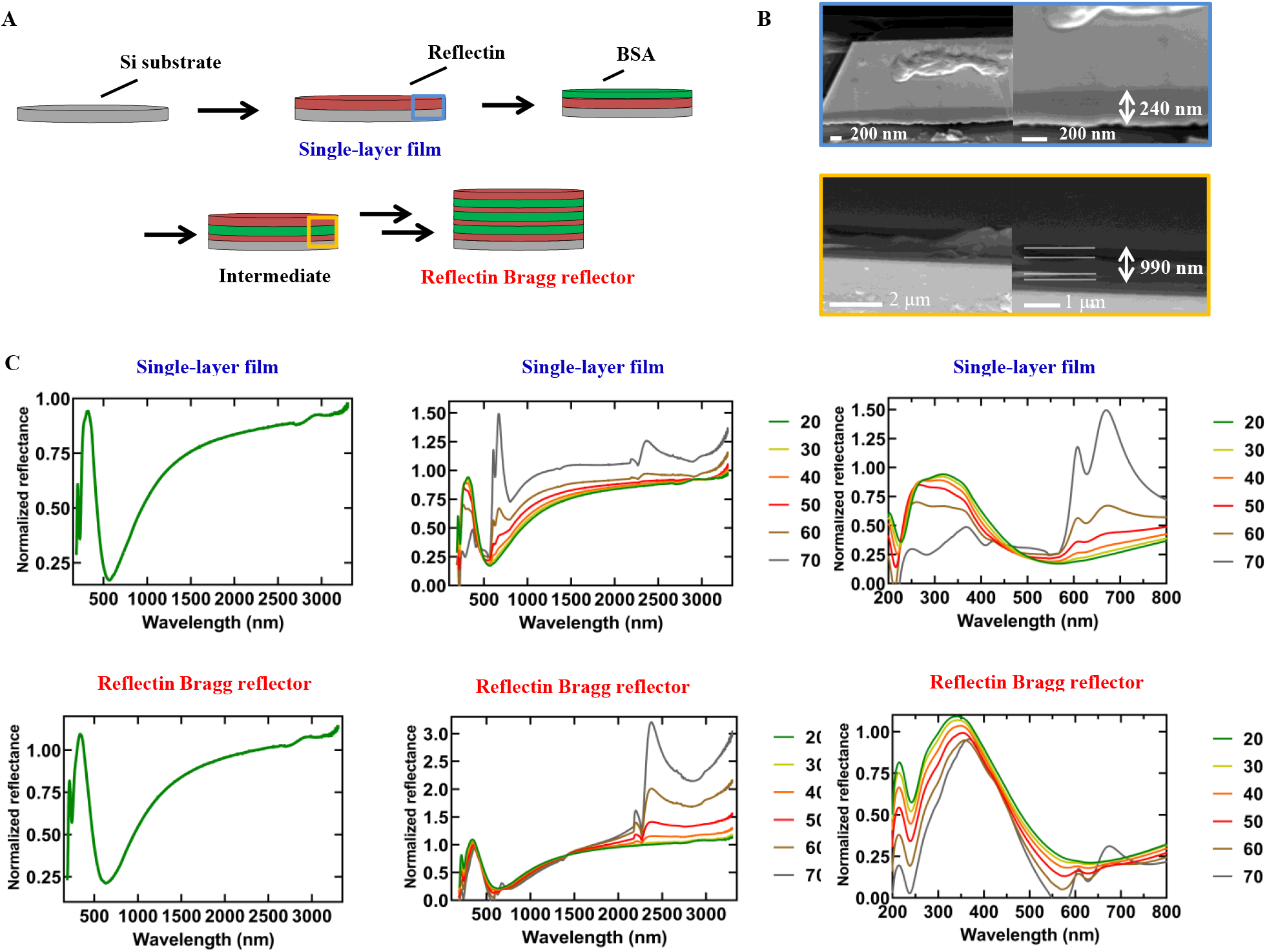
A) Schematic figure showing the fabrication of reflectin single- and multilayer thin-films B) Cross-sectional SEM of single layer film (top) and tri-layered intermediate (bottom). C) UV-Vis-NIR reflectance spectra of reflectin single-layer films (top) and reflectin Bragg reflectors (Bottom). Left column; full spectra at 20°, middle column; full spectra at 20-70°, right column; UV-Vis spectra at 20-70°.

Designing multilayer films provided the opportunity to investigate incorporating other proteins into reflectin multilayer films. Of particular interest is the use of novel methods to control the thickness and therefore optical properties of reflectin materials. Towards this purpose, the phytochrome family of photoreceptor proteins is an attractive target as they undergo a reversible light-induced conversion between two different states, which involves a significant conformational change in the protein (Figure S7). ^[28,29]^ In solution, after being exposed to 625 nm light, the UV spectra of this phytochrome was found to contain a peak around 750 nm, which corresponds to an ‘open’ state of the protein. After exposure to 780 nm light, the peak shifted to ∼710 nm, corresponding to a ‘closed’ state of the protein (Figure 3A). A reflectin/phytochrome device was then designed and subsequently fabricated by spin coating a layer of phytochrome onto a clean Si wafer followed by a layer of reflectin, with the presence of discrete layers confirmed using cross-sectional SEM (Figure 3B). The optical properties were then characterized in the visible region, revealing peak reflectance around 600 nm. It was then possible to trigger photoisomerism of the coated phytochrome by shining 625/780 nm light onto the surface of the film in the presence of water vapor (Figure 3C). Continuous exposure to 625 nm light in the presence of water vapor resulted in a red-shift to ∼610 nm, while exposure to 780 nm light in the presence of water vapor resulted in a blue-shift to ∼598 nm. It was also possible to cycle multiple times between the two states by switching between red and far-red light, representing a novel, reversible method of controlling the optical properties of reflectin-based materials, which may lead to a new class of photoinducible reflectin-based camouflage devices.

**Figure 3.**
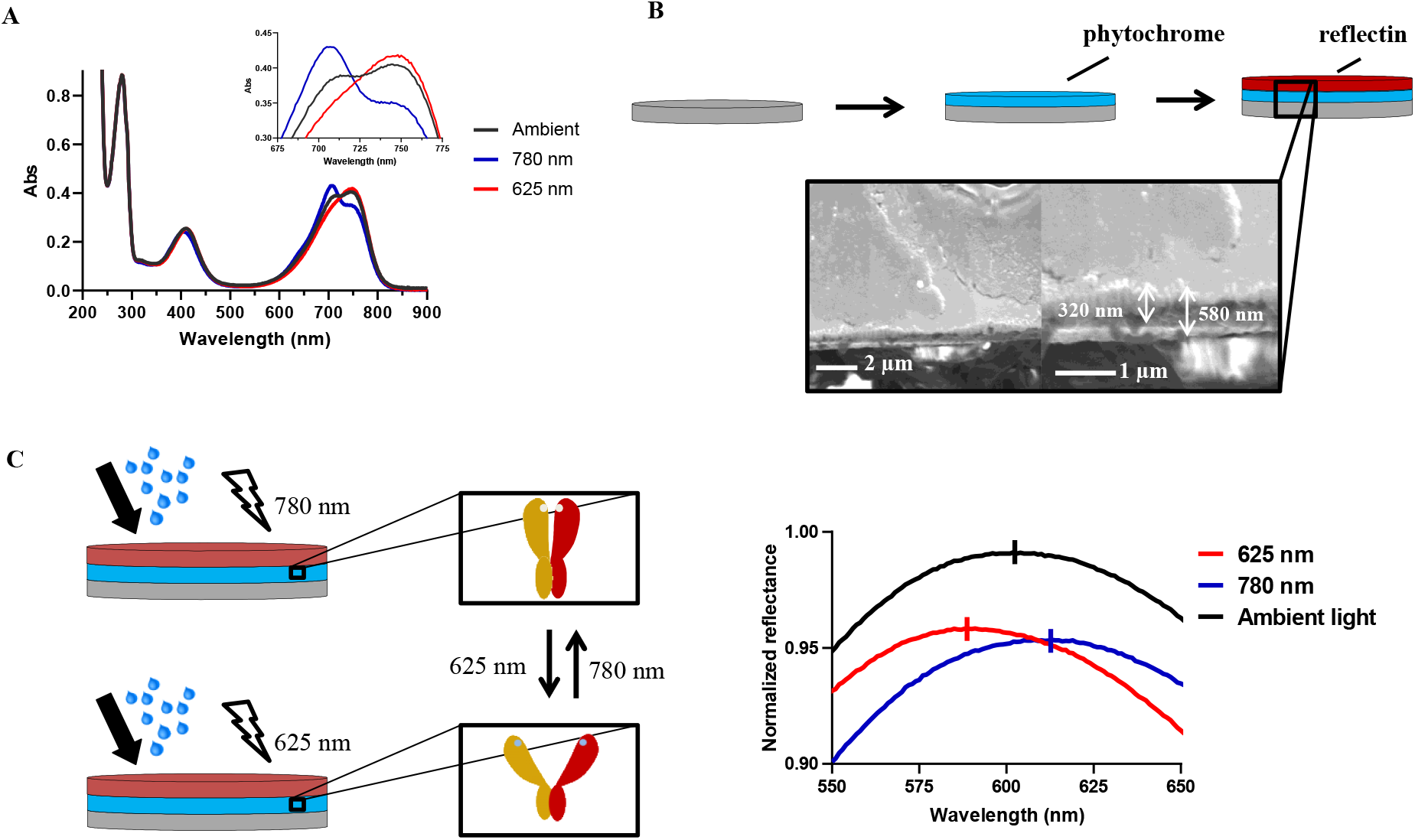
A) UV/Vis spectra of phytochrome Cpar GPS1 in H_2_O after being exposed to ambient, red (625 nm), and far-red (780 nm) light. B) Schematic figure showing the fabrication of reflectin-phytochrome device and corresponding cross-sectional SEM image. C) Schematic figure of reflectin-phytochrome device switching between ‘open’ and ‘closed’ states along with the corresponding reflectance spectra.

In conclusion, characterizing reflectin-based single layer films has revealed previously unreported optical properties conserved across all samples, such as UV reflectivity and broad reflectance in the NIR/short-wave IR regions extending to 3300 nm. The angle-dependent-reflectivity of single-layer films has been determined, revealing significant spectral shifts associated with changes to the angle of incidence. Moving towards a more biomimetic configuration led to reduced angle-dependence while maintaining the spectral features of single-layer films, representing an important step towards developing angle-independent reflectin-based camouflage technologies. Finally, the design and fabrication of a novel reflectin-phytochrome device, controlled using visible light-induced isomerism, was described, adding to the catalogue of methods to control reflectin-based materials optical properties post-fabrication.

## EXPERIMENTAL SECTION

### Protein expression and purification

The genes encoding reflectins had been digested by NdeI and XhoI and ligated into either a pETM11(+) vector (containing a kanamycin resistance gene) or a PMA-T vector (containing an ampicillin resistance gene) digested by the same restriction enzymes, yielding pETM11_XXRefYY/PMa-T_XXRefYY, where XX corresponds to the origin of the reflectin gene, and YY corresponds to the reflectin isoform. Vectors (pETM11_ESRef1a, pETM11_ESRef2D, pETM11_DORefA2, pETM11_ESRef2A, and pETM11_SORef1, were transformed into BL21(DE3) cells (Novagen) and expressed at 37 °C using Overnight Express Instant Terrific Broth (TB) or LB media (Formedium) supplemented with 30 μg mL^−1^ kanamycin/ 50 μg mL^−1^ ampicillin. All isoforms were completely insoluble when expressed in E.coli and were sequestered within inclusion bodies. Inclusion bodies were resuspended in buffer A (20 mM Tris HCl, 100 mM NaCl), sonicated (5 × 15 seconds, 45 second intervals, 40%), and centrifuged (21000 g, 30 mins). The pellet was then resuspended in wash buffer (20 mM Tris HCl, 5 mM EDTA, 5 mM DTT, 2M Urea) supplemented with 2% Triton and centrifuged (21000 g, 30 mins). This triton wash was then repeated, followed by centrifugation (21000 g, 30 mins) and pellet resuspension in wash buffer. After a final centrifugation step (21000 g, 30 min) the pellet was solubilized in denaturing buffer (6 M guanidine hydrochloride), adjusted to pH 8 using NaOH, and left stirring overnight. The sample was then filtered through 5, 0.4, and 0.2 μm sterile filters, clarified by centrifugation (48000 g, 60 mins), and purified using HPLC on an Agilent 1260 Infinity system using a reverse phase C18 column. Elution conditions: 95% Buffer A:5% Buffer B to 5% Buffer A:95% Buffer B at a flow rate of 5-25 mL min^−1^ over 20 minutes (Buffer A: 99.9% water, 0.1% trifluoroacetic acid; Buffer B: 99.9% acetonitrile, 0.1% trifluoroacetic acid). The pure fractions were pooled, flash frozen in liquid nitrogen, and lyophilized. Protein purity was assessed by SDS-PAGE on 4–12% Bis–Tris precast gels (Bio-Rad, USA).

### E. coli transformation for plasmid preparations

Plasmid DNA (1 μL) was mixed with 25-50 μL of competent cells and the mixture placed on ice for 30 min. The sample was heat-shocked at 42 °C for 10-30 seconds and placed on ice for 5 min. SOC medium (500-950 μL) was added at room temperature and cells incubated at 37 °C for 45-60 min. The cells were spread onto a kanamycin (50 μgml^−1^) or ampicillin (30 μgml^−1^) antibiotic-selective plate and incubated overnight at 37 °C. For DNA amplification the transformations were done using DH5α competent cells, while for protein expression BL21(DE3) was used. A single colony from each plate was used to inoculate 10 mL of LB medium supplemented with kanamycin (30 μg ml^−1^) or ampicillin (50 μg ml^−1^). The starter cultures were incubated at 37 °C @ 180 rpm overnight. 500 μl of the culture was mixed with 500 μl of a sterile 50% glycerol solution, flash-frozen in liquid N2, and stored at −80°C. For plasmid preparations, QIAGEN spin miniprep kit was used following the recommended protocol. DNA concentrations were determined using the NanoDrop™ 2000.

### UV/Vis spectroscopy

Samples were dissolved in MQ H_2_O at 0.5 mg/mL and incubated overnight. UV/vis spectra were collected using a Cary 60 UV-Vis spectrophotometer (Agilent technologies) from 200-900 nm.

### Atomic-force microscopy

AFM imaging was carried out in contact mode using an Asylum Research MFP-3D (Oxford Instruments, High Wycombe, UK) atomic force microscope using a NuSense Scout 350 cantilever (NuNano, Bristol, UK). AFM data was analysed with the Gwyddion software package, http://gwyddion.net/.

### Cross-sectional SEM

SEM was carried out on the cross-section of reflectin-based films using an FEI Quanta 250 microscope with accelerating voltage of 5 to 8 kV. Samples were sputter-coated with 5 nm Au/Pd to enhance electrical conductivity.

### Reflectance spectrophotometry

The reflectance spectra of 1% reflectin thin-films were collected at room temperature using an Agilent Cary 5000 series UV-VIS-NIR spectrophotometer with a variable angle specular reflectance accessory (VARSA). Spectra were collected between 175-3300 nm with a scan rate of 600 nm min^−1^ (1 nm data interval, average time 0.1 s). The angle of incidence was varied between 20-70 degrees (10 degree intervals). For reflectin/phytochrome samples, reflectance was collected between 400-800 nm.

### Spin-coating

Reflectin samples were dissolved in hexafluoroisopropanol (HFIP) at 1% (w/w). Silicon wafer substrates were cleaned with piranha solution (3:1 of sulfuric acid and 30% hydrogen peroxide, hazardous solution) for 1 h, rinsed with HPLC-grade water, polished using lens tissue, rinsed with IPA, and spun at 1500rpm for 1 minute to dry. 100 μL reflectin sample was deposited onto the substrates before spin coating at 1500 rpm for 60 s. Films were allowed to dry in a fume hood at room temperature. A2/BSA multilayer films were fabricated by first spin-coating a 1% (w/w) sample of DORefA2 in HFIP onto a silicon wafer, and once dry, further spin coating a 1% (w/w) sample of BSA in HFIP. This process was then repeated 2.5 times to yield a multilayer film with seven alternating layers; A:B:A:B:A:B:A, where A is DORefA2 and B is BSA. Reflectin/phytochrome thin-films were fabricated by first spin-coating phytochrome (5 mg/mL in HFIP/H2O) onto a Si wafer, followed by spin coating DOrefA2 (1 mg/mL in HFIP).

### Photoisomerism

Reflectin/phytochrome samples were continuously exposed to LED light (625 or 780 nm) for 3 minutes in a dark room while water vapour was applied to the surface every 30 seconds.

## SUPPORTING INFORMATION

Supporting Information is available online or from the author.

## ACKNOWLEDGEMENTS

This work was supported by the Future Biomanufacturing Research Hub (grant EP/S01778X/1), funded by the Engineering and Physical Sciences Research Council (EPSRC) and Biotechnology and Biological Sciences Research Council (BBSRC) as part of UK Research and Innovation. This work was also supported by the UK Defence Science and Technology Laboratory (PhD studentship to E. W-M). The authors thank the University of Manchester BioImaging (BioAFM) Facility for assistance with AFM.

## ToC Figure

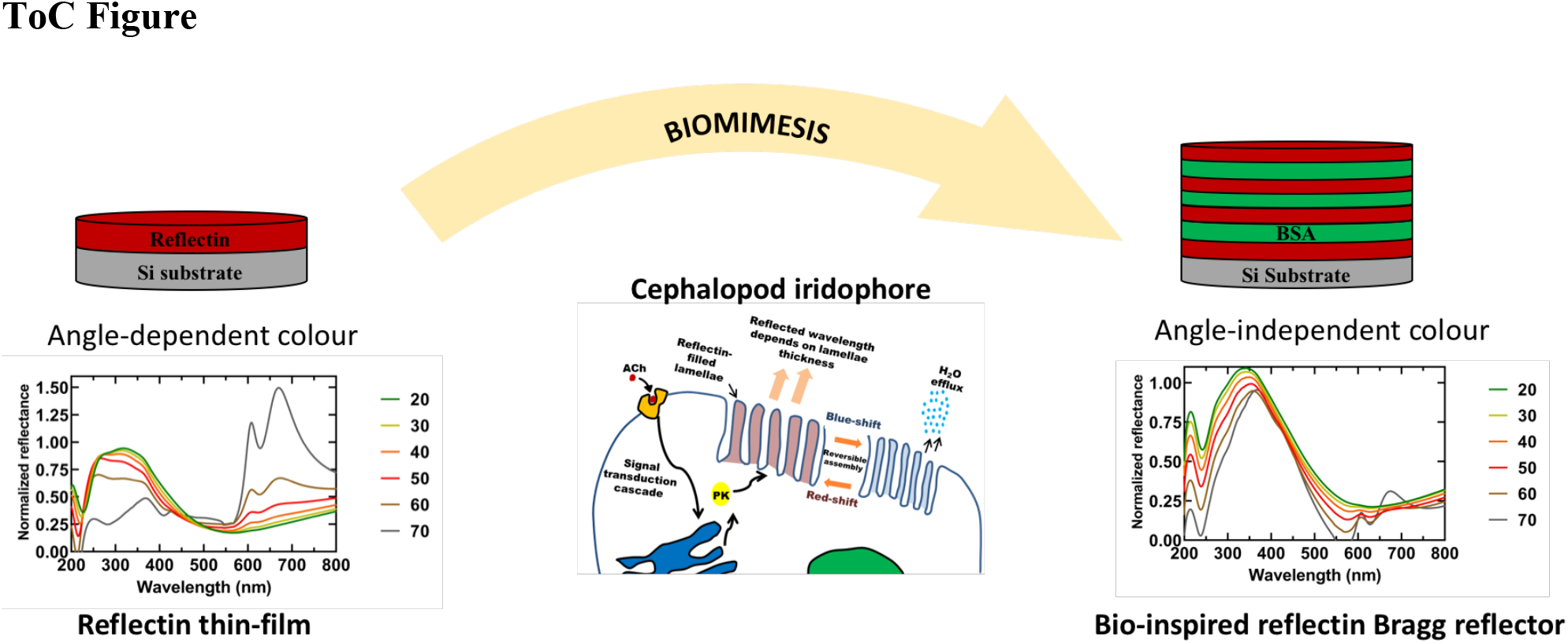
**ToC text:** The angular-dependent optical properties of reflectin single layer thin-films is minimized by taking more direct inspiration from natural cephalopod iridophores, leading to the design and fabrication of a bio-inspired reflectin Bragg reflector. Our design was shown to conserve the optical properties of single-layer films while significantly reducing the effect of shifting angle of incidence on specular reflectance.

