## Supplementary data for "Design and fabrication of recombinant reflectin-based Bragg reflectors: bio-design engineering and photoisomerism induced wavelength modulation"

### Supporting Information

**Reflectin 1A [Euprymna scolopes] AAQ21389:**
MNRFMNRYRPMFNNMYSNMYRGRYRGM**MEPMSRMTMDFQGRYMDSQGRM**VDPRYYDYYGRFNDYD
RYYGRSMFNYGWMMDGDRYNRYNRW**MDYPERYMDMSGYQMDMSGRWMDMQGRHC**NPYSQWMM
YNYNRHGYYPNYSYGRHMFYPERW**MDMSNYSMDMYGRYMDRWGRY**CNPFSQYMNYYGRYWNYPGYN
NYYYSRNMYYPERY**FDMSNWQMDMQGRWMDNQGRY**CSPYWNNWYGRHMYYPYQNNYFYGRYDYPG
**MDYSNYQMDMQGRYMDQYGMN**DYYY

**Reflectin 2A [Euprymna scolopes] AAQ21392:**
MNRYMTRFRNFYGNMYRGRYRGM**MEPMSRMTMDFQGRYMDSQGRM**VDPRYYDYYGRYNDYDRYYGRS
MFNYGWMMDGDRYNRYNRW**MDFPERYMDMSGYQMDMYGRWMDMQGRH**CNPYSQMMYNYNRHG
YYPNYSYGRHMFYPERW**MDMSNYSMDMYGRYMDRWGRY**CNPFYQFYNHWNRYGNYPGYYNYYYMYYP
ERY**FDMSNWQMDMQGRWMDMQGRY**CSPYWYNWYGRHMYYPYQNYYWYGRYDYPG**MDYSNWQMD**
**MQGRWMDMQGRYMDYPYNY**YNWY

**Sequence 46 (reflectin 2D) [Euprymna Scolopes] US 7314735 ABZ24324:**
MNRYMNRFRNFYGNMYRGRYRGM**MEPMSRMTMDFQGRYMDSQGRM**VDPRYYDYYGRFNDYDRYYGR
SMFNYGWMMDGDRYNRYNRW**MDFPERYMDMSGYQMDMYGRWMDMQGRH**CNPYSQWMMYNYN
RHGYYPNYSYGRHMFYPERW**MDMSNYSMDMYGRYMDRWGRY**CNPFYQFYNHWNRYGNYPGYYSYYYM
YYPERY**FDMSNWQMDMQGRWMDMQGRY**CSPYWYNWYGRHMYYPYQNYYWYGRYDYPG**MDYSNWQ**
**MDMQGRWMDMQGRYMDYPYNY**YNWNH

**Reflectin 1 [Sepia officinalis] CCG28037:**
MNRYMMRNRPMYGNMYRTGKKYRGV**MEPMSRMTMDFQGRYMDSQGRM**VDPRHNDYYGRWNDYDR
YYGRSMFNYGPHMDGHQHGGW**MDFPERWMDMSNYQMDMQGRWMDMQGRH**CQPFNQWGYNRH
GNYPSSYYGRN**MFYPERWMDMSNWQMDTQGRWMDMQGRY**GSPFNQWGYNRHGYYPGSSYGRNMY
HPERW**MDMSNYQMDMQGRWMDMHGRH**VNPFSHSMHGRNWSYPYYNYYSSRH**MDYPERNMDMSN**
**WQMDMQGRWMDMQGRHMDPSWSN**MHDNHNYWF

**Reflectin A2 [Doryteuthis opalescens] KF661516:**
MNRYMMRHRPMYSNMYRTGRKYRGV**MEPMSRMTMDFQGRYMDSQGRM**VDPRYYEYGRCHDYDRYNG
RSMFNNGPYMDGQRYGGW**MDFPERYMDMSGYQMDMHGRWMDSQGRY**CNPMGHSWSNRQGYYPGS
NYGRNMFNPERY**MDMSGYQMDMQGRWMDMGGRH**VNPFSHSMYGRNMFNPSYFSNRH**MDNPERYM**
**DMSGYQMDMQGRWMDTQGRYMDPSMSN**MYDNYNYWY

**Figure S1** Name, origin, UniProt ID, and amino-acid sequence of all reflectin isoforms used in this study. Repeating motif regions are highlighted bold.

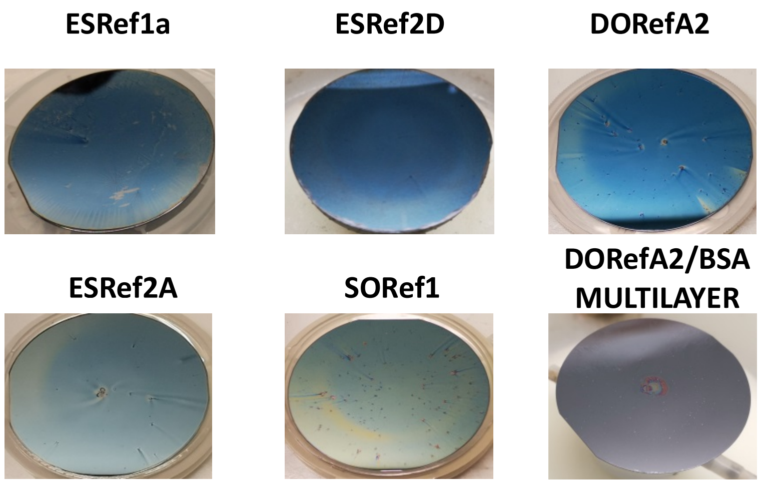


**Figure S2** Optical images of reflectin thin-films following fabrication by spin coating 1% w/w reflectin in HFIP onto a clean Si wafer. Camera images were taken following drying. DORefA2/BSA multilayer films were fabricated by spin coating reflectin and BSA sequentially.

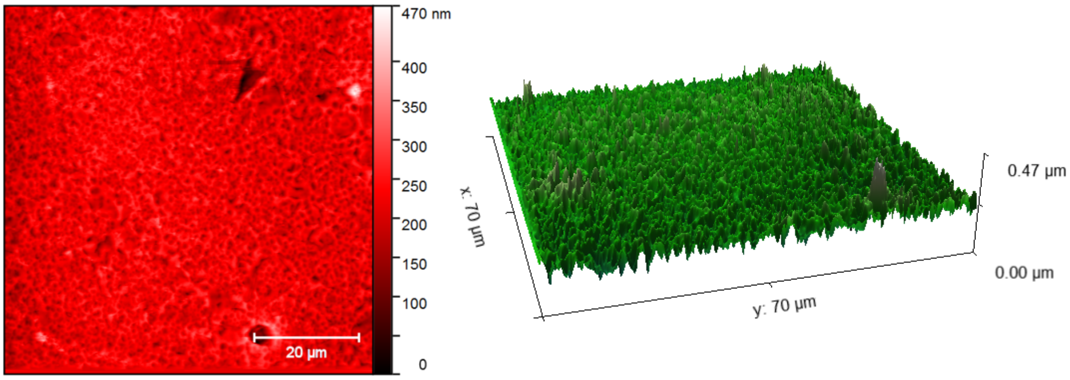


**Figure S3** Representative atomic force microscopy (AFM) images of a DORefA2 thin-film fabricated by spin coating 1% w/w DORefA2 in HFIP onto a clean Si wafer. AFM data was processed using the Gwyddion software package, http://gwyddion.net/.

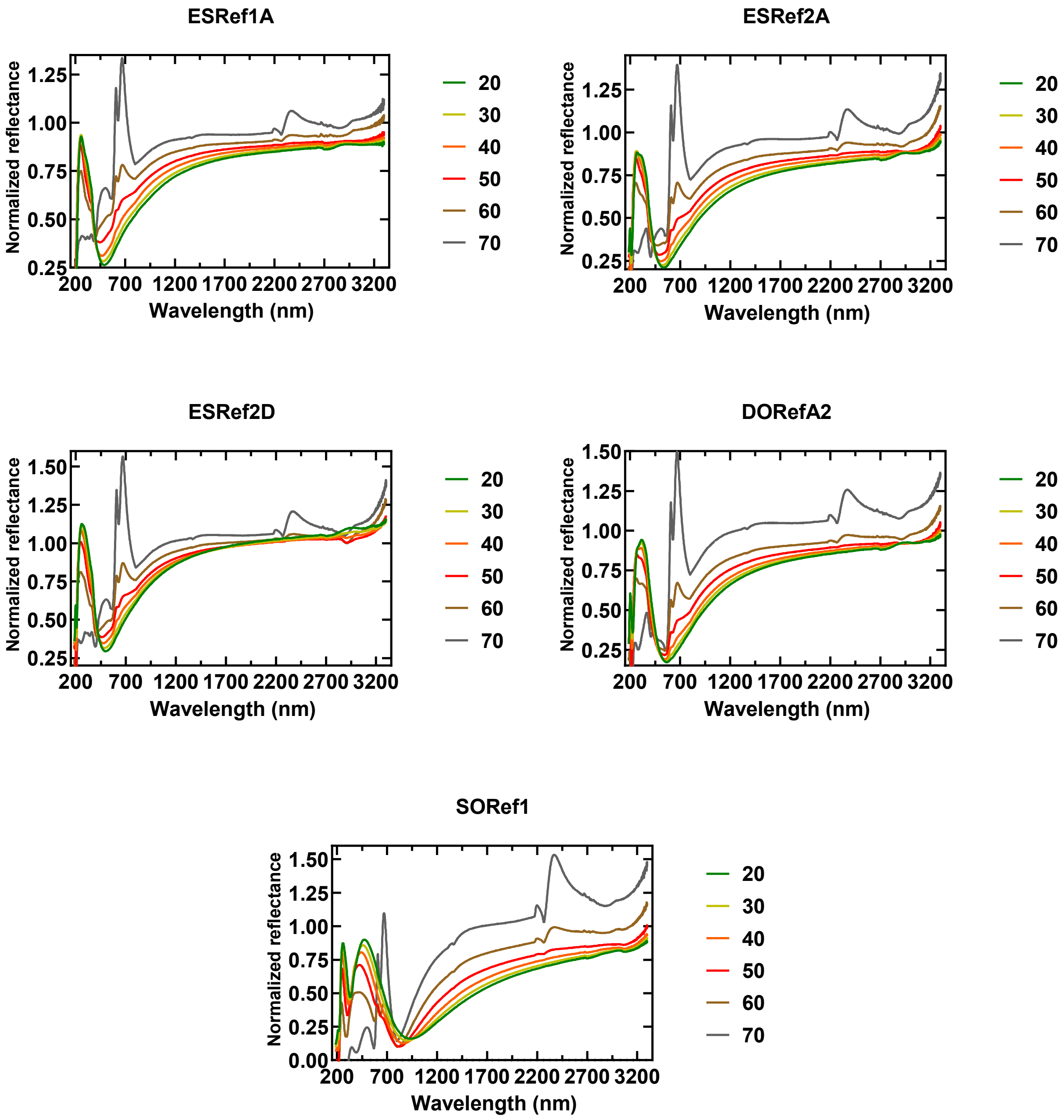


**Figure S4** UV-Vis-NIR reflectance spectra (185-3300 nm) of reflectin-based single-layer films fabricated by spin coating 1% w/w re flectin in HFIP onto clean Si wafers. The angle of incidence was varied between 20-70 degrees (10 degree intervals).

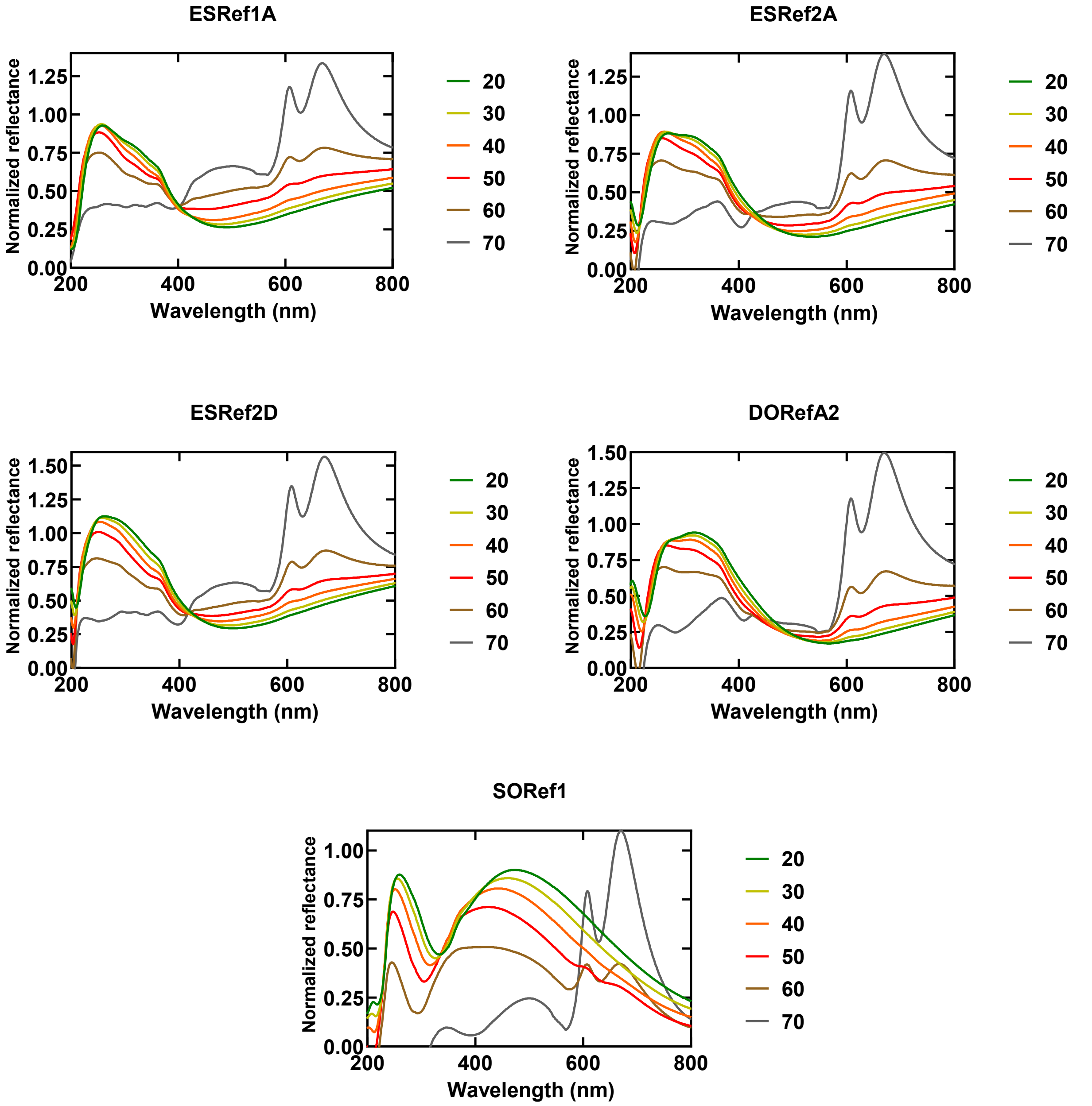


**Figure S5** UV-Vis reflectance spectra (200-800 nm) of reflectin-based single-layer films fabricated by spin coating 1% w/w re flectin in HFIP onto clean Si wafers. The angle of incidence was varied between 20-70 degrees (10 degree intervals).

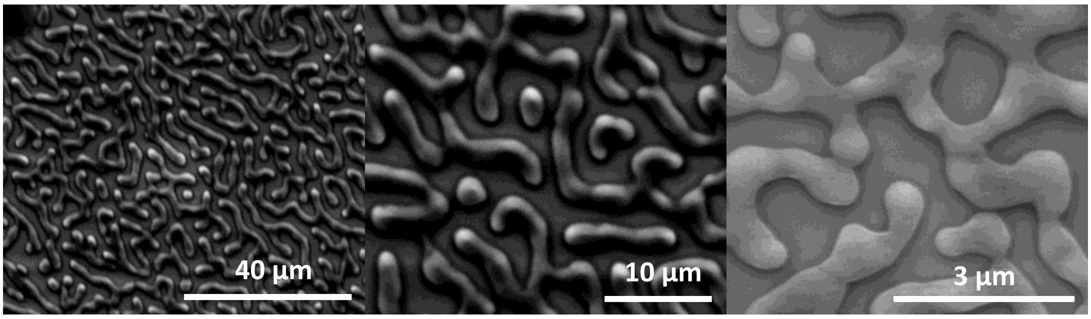


**Figure S6** Representative SEM micrographs of the surface of a reflectin thin-film fabricated by sping coating 1% w/w reflectin in HFIP onto a clean Si wafer.

**Agrobacterium fabrum PHY1:**
MSSHTPKLDSCGAEPIHIPGAIQEHGALLVLSAREFSVVQASDNLANYIGVDLPIGAVATEA
NLPFISVLSAWYSGEESNFRYAWAEKKLDVSAHRSGTLVILEVEKAGVGESAEKLMGELTS
LAKYLNSAPSLEDALFRTAQLVSSISGHDRTLIYDFGLDWSGHVVAEAGSGALPSYLGLRF
PAGDIPPQARQLYTINRLRMIPDVDYKPVPIRPEVNAETGAVLDMSFSQLRSVSPVHLEYMR
NMGTAASMSVSIVVNGALWGLIACHHATPHSVSLAVREACDFAAQLLSMRIAMEQSSQD
ASRRVELGHIQARLLKGMAAAEKWVDGLLGGEGEREDLLKQVGADGAALVLGDDYELV
GNTPSREQVEELILWLGEREIADVFATDNLAGNYPTAAAYASVASGIIAMRVSELHGSWLI
WFRPEVIKTVRWGGDPHKTVQESGRIHPRKSFEIWKEQLRNTSFPWSEPELAAARELRGAII
GIVLRKTEEMADLTRELQRTNKELEAFSYSVSHDLRAPFRHIVGFAQLLRERSDALDEKSL
HYLQMISEAALGAGRLVDDLLNFSQLGRTQLTLKPVDMQKVVSEVRRSLSHAVSDRQIEW
RIGALPVIFGDPTLLRQVWYNLIENAIKYSSREPVSIITISAVETEDDVTYSVEDNGVGFDMA
YYNKLFGVFQRLQRVEDFEGTGIGLALVRRIVERHHGLVGAEGTVGEGATFSFTLPVTKVE
EEKIA

**Figure S7** Amino acid sequence of phytochrome 1 from Agrobacterium fabrum used in this study.
